# REM sleep quality is associated with balanced tonic activity of the locus coeruleus during wakefulness

**DOI:** 10.1101/2024.07.12.603275

**Authors:** Nasrin Mortazavi, Puneet Talwar, Ekaterina Koshmanova, Roya Sharifpour, Elise Beckers, Alexandre Berger, Islay Campbell, Ilenia Paparella, Fermin Balda, Ismael Dardour Hamzaoui, Christian Berthomier, Christine Bastin, Christophe Phillips, Pierre Maquet, Fabienne Collette, Mikhail Zubkov, Laurent Lamalle, Gilles Vandewalle

## Abstract

**Background:** Animal studies established that the locus coeruleus (LC) plays important roles in sleep and wakefulness regulation. Whether it contributes to sleep variability in humans is not yet established. Here, we investigated if the *in vivo* activity of the LC is related to the variability in the quality of Rapid Eye Movement (REM) sleep.

**Methods:** We assessed the LC activity of 34 healthy younger (∼22y) and 18 older (∼61y) individuals engaged in bottom-up and top-down cognitive tasks using 7-Tesla functional Magnetic Resonance Imaging (fMRI). We further recorded their sleep electroencephalogram (EEG) to evaluate associations between LC fMRI measures and REM sleep EEG metrics.

**Results:** Theta oscillation energy during REM sleep was positively associated with LC response in the top-down task. In contrast, REM sleep theta energy was negatively associated with LC activity in older individuals during the bottom-up task. Importantly, sigma oscillations power immediately preceding a REM sleep episode was positively associated with LC activity in the top-down task.

**Conclusions:** LC activity during wakefulness was related to REM sleep intensity and to a transient EEG change preceding REM sleep, a feature causally related to LC activity in animal studies. The associations depend on the cognitive task, suggesting that a balanced level of LC tonic activity during wakefulness is required for optimal expression of REM sleep. The findings may have implications for the high prevalence of sleep complaints reported in aging and for disorders such as insomnia, Alzheimer’s, and Parkinson’s disease, for which the LC may play pivotal roles through sleep.

## Background

Up to 35% of the general population report unsatisfying sleep[1] and one out of two adults aged over 50y complains about sleep or daytime sleepiness.[2] Insomnia is the obvious extreme form of these complaints and constitutes the second most prevalent psychiatric disorder.[3] Beyond the behaviors that do not favor appropriate sleep, some people are more vulnerable to poor sleep or more at risk of developing insomnia.[4,5] The biological origin of sleep variability is however not fully established.[6]

The locus coeruleus (LC) is a small nucleus of the dorsal pons – cylinder-like shape of approximately 15 mm with ∼2.5 mm diameter.[7] It is responsible for producing norepinephrine (NE) as a part of the ascending arousal system and projects to the neocortex, hippocampus, amygdala, thalamus, and cerebellum.[8] It plays a crucial role in the control of behavioral states, including alertness and attention,[9] as well as in the regulation of sleep.[6] LC-NE tone must decrease to allow sleep onset, while LC activity dynamically changes during sleep to shape the alternation between Rapid Eye Movement (REM) and non-REM sleep and some of the microstructure elements of sleep.[10,11] A lot of our understanding of the role of the LC in sleep regulation arises from experiments conducted in rodents. Therefore translation to diurnal human beings, with a more developed cortex, is not straightforward[6]. In addition, imaging the LC *in vivo* remains difficult because of its deep location and small size, as well as because of the difficulty to image human sleeping brains. Recent advances in neuroimaging techniques have lifted part of these limitations.[12] For instance, degeneration of the LC was recently suggested to contribute to alteration in rest-activity patterns.[12] We further reported that a higher response of the LC to a salience detection task during wakefulness, which presumably reflects in part LC activity during sleep, was associated with lower intensity of REM sleep in healthy older individuals.[13] REM sleep intensity does not however solely depend on LC activity, and no association with sleep features shown to be under the direct control of the LC in animal were reported in the study.

LC neurons follow phasic and tonic activity patterns, which lead to distinct release of NE.[14] Phasic activity typically occurs in response to transient salient stimulation, while the tonic mode is mostly required for sustained processes. The dynamics of the tonic mode is considered to follow an inverted U-shape where low LC tonic discharge is associated with low arousal and poor performance, moderate LC tonic activity allowing pronounced phasic activity leads to optimal arousal and performance, and high tonic activity hinders phasic activity and leads to anxiety-like behavior and poorer performance .[15] Likewise, during sleep the LC undergoes periods of higher or lower tonic activity.[10,11]

How the balance between phasic and tonic modes of activity of the LC during wakefulness may be related to REM sleep is not known. In the present study, we used Ultra-High-Field (UHF) 7- Tesla (7T) Magnetic Resonance Imaging (MRI) to extract *in vivo* the LC activity in healthy younger and older adults during two cognitive tasks that rely on different tonic tones of the LC. We related the LC activity during both tasks to electrophysiology metrics of REM sleep. We hypothesized that the LC activity in both tasks would be mainly associated with REM sleep intensity. Moreover, we expected that the associations would differ between two tasks, and would be more pronounced in older individuals. To test for associations with sleep features under direct control of the LC, we further tested the association between LC activity and the power of the sigma band of the EEG that preceded REM sleep episodes, because it is causally related to NE tone in rodents[16].

## Methods

This study was approved by the faculty-hospital ethics committee of ULiège. All participants provided written informed consent and received financial compensation. The study is part of a larger project that has led to previous publications.[13,17] Part of the results dealing with one cognitive task were reported in one of these publications[13] and are included for comparisons purposes and to conduct additional analyses. Most of the methods were described in details in[13]. More details are also available in the **Supplementary Methods**.

### Participants

Fifty-two healthy participants including 34 healthy young (22.2±3.1y, 28 women) and 18 late middle-aged (60.9±5.4y, 14 women) individuals, completed the study **(Table 1)**. The exclusion criteria were as follows: history of major neurologic/psychiatric diseases or stroke; recent history of depression/anxiety; sleep disorders; medication affecting the central nervous system; smoking, excessive alcohol (>14 units/week) or caffeine (>5 cups/day) consumption; night shift work in the past 6 months; BMI ≤18 and ≥29 (for older individuals) and ≥25 (for younger individuals). All older participants had to show normal performance on the Mattis Dementia Rating Scale (score > 130/144).[18]

**Table 1.** Characteristics of the study sample.

|  | Young individuals<br>(n = 34) |  |  |  | Late middle-aged individuals<br>(n = 18) |  |  |  | P-value |
| --- | --- | --- | --- | --- | --- | --- | --- | --- | --- |
|  | Mean | SD | Min | Max | Mean | SD | Min | Max |  |
| Age (years) | 22.20 | 3.17 | 18.00 | 29.00 | 60.88 | 5.41 | 53.00 | 70.00 | <.0001 |
| BMI (kg/m<sup>2</sup>) | 21.90 | 2.74 | 17.20 | 28.40 | 24.75 | 3.40 | 19.40 | 30.90 | **0.0047** |
| Education (years) | 14.46 | 2.31 | 12.00 | 20.00 | 14.61 | 2.70 | 9.00 | 19.00 | 0.85 |
| Depression level (BDI) | 6.64 | 4.17 | 0 | 20.00 | 5.33 | 4.01 | 0 | 14.00 | 0.27 |
| Anxiety level (BAI) | 4.00 | 2.92 | 0 | 11.00 | 3.05 | 3.18 | 0 | 9.00 | 0.30 |
| TIV (cm<sup>3</sup>) | 1402.1 | 136.6 | 1203.3 | 1946.7 | 1420.5 | 125.5 | 1167.6 | 1600.1 | 0.627 |
| Daytime sleepiness (ESS) | 6.93 | 3.90 | 0 | 14.00 | 5.77 | 3.76 | 0 | 13.00 | 0.30 |
| Sex (F–M) | 28 F – 6 M |  |  |  | 12 F – 6 M |  |  |  | 0.20 |
| Habitual Subjective sleep quality (PSQI) | 4.79 | 2.18 | 1.00 | 10.00 | 3.83 | 2.22 | 0 | 8.00 | 0.14 |
| TST (min) | 454.80 | 30.97 | 379.50 | 499.50 | 394.63 | 48.16 | 282.50 | 495.50 | <.0001 |
| Theta power in REMS (μV<sup>2</sup>) | 7281.5 | 5061.7 | 1162.4 | 22346.4 | 4789.5 | 3508.2 | 765.3 | 13051.8 | **0.0449** |
| REMS percentage (%) | 26.63 | 4.41 | 19.93 | 36.45 | 21.52 | 5.50 | 13.11 | 35.39 | **0.002** |
| REMS latency (min) | 103.0 | 47.834 | 0 | 213.5 | 81.944 | 56.806 | 38.000 | 295.5 | 0.190 |
| Sigma power prior to REMS (μV<sup>2</sup>) | 15.7615 | 8.8957 | 5.0656 | 44.356 | 16.0544 | 6.795 | 3.654 | 27.856 | 0.896 |
| SWE (μV<sup>2</sup>) | 175531 | 171295 | 14259.2 | 555453 | 126388 | 109290 | 23293.3 | 470116 | 0.2146 |

The p-values shown in the table correspond to two-sample t-tests except for sex that were compared
using a Chi-square test. BMI: Body Mass Index; BDI: Beck Depression Inventory; BAI: Beck Anxiety
Inventory; TIV: Total Intracranial Volume; PSQI: Pittsburgh Sleep Quality Index; ESS: Epworth Sleepiness
Scale; TST: Total Sleep Time; SWE: slow wave energy; REMS: rapid eye movement sleep. See
supplementary method for references of the questionnaires.

### Protocol

Participants’ sleep was recorded in the lab twice. First, participants completed a night of sleep under polysomnography (called habituation night) to screen and exclude for sleep abnormalities (apnea hourly index and periodic leg movement >15; no parasomnia or REM behavioral disorder). All participants further completed a whole-brain structural MRI (sMRI) acquisition and a specific acquisition centered on the LC. Participants were then requested to sleep regularly for 7 days before the baseline night (±30min from their sleep schedule) based on their preferred schedule (compliance was verified using sleep diaries and wrist actigraphy - Actiwatch and AX3, AXIVITY LTD, Newcastle, UK). For the baseline night, participants first remained awake for 3h under dim light (<10 lux) then their habitual sleep was recorded in darkness (N7000 amplifier, EMBLA, Natus, Middleton, WI). Approximately 3h after wake-up time under dim light (<10 lux), participants completed a functional MRI (fMRI) session that included 3 tasks (**Figure 1A**). This paper is centered on the analyses of the first two tasks, the perceptual rivalry and auditory salience detection tasks.

**Figure 1.**
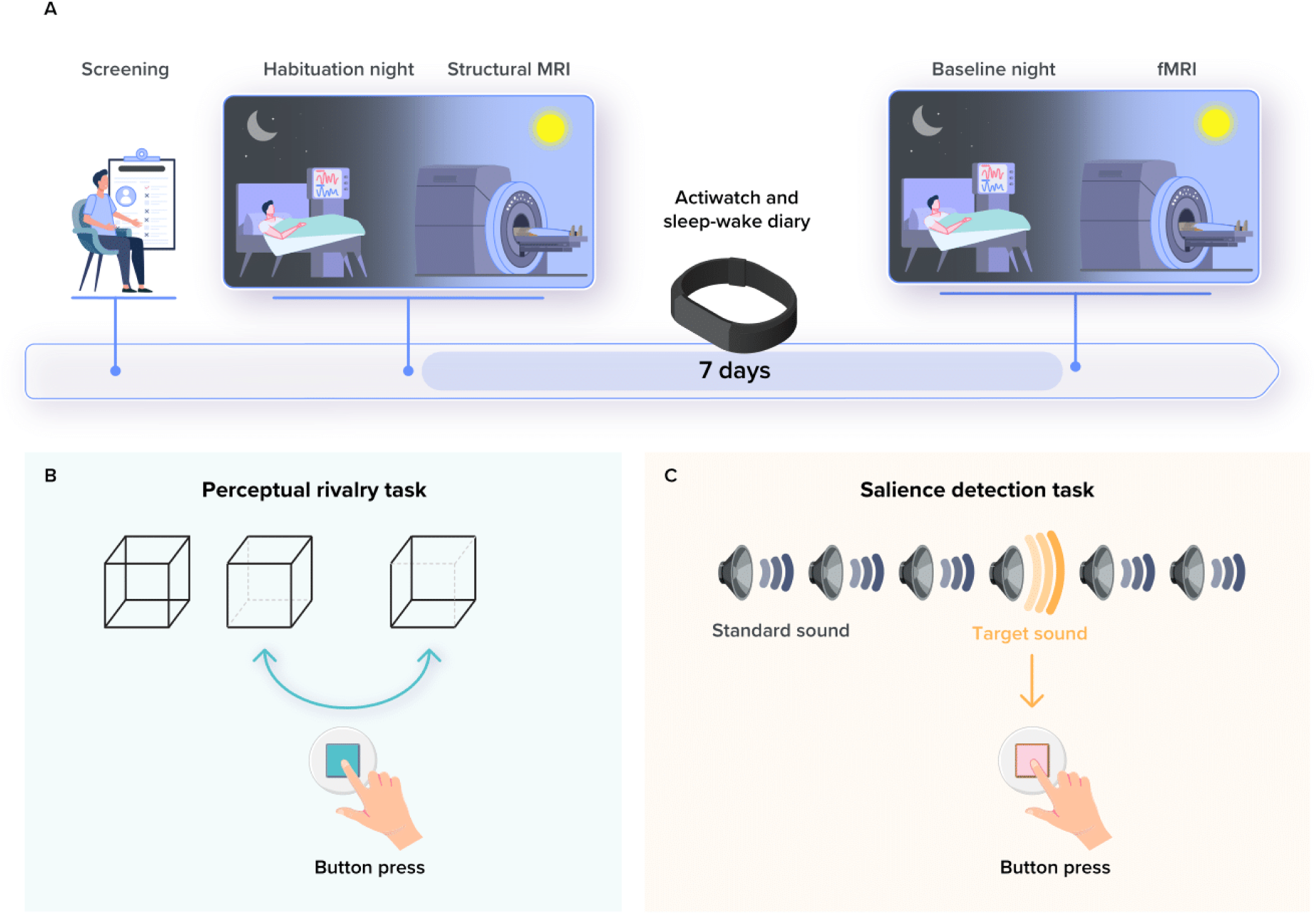
Overview of the study protocol. **(A)** After screening, participants completed an in-lab screening and habituation night under polysomnography to minimize the effect of the novel environment for the subsequent baseline night and to exclude volunteers with sleep disorders. They further completed a structural 7T MRI session including a whole-brain structural MRI and a LC-specific sequence. The latter was used to create individual LC masks in each participant’s brain space. After 7 nights of regular sleep-wake time at home, which was confirmed by actigraphy data and/or sleep-wake diary, participants came to the lab three hours before their sleep time and were maintained in dim light (<10 lux) until sleep time. Participants’ habitual baseline sleep were recorded overnight in-lab under EEG to extract our main sleep features of interest. All participants underwent an fMRI session approximately 3h after wake-up time (following ≥45min in dim light - < 10lux), during which they completed the visual perceptual rivalry task and the auditory salience detection task. **(B)** The Visual perceptual rivalry task consisted of watching a 3D Necker cube, which can be perceived in two different orientations (blue arrow), for 10 blocks of 1min separated by 10s of screen-center cross fixation (total duration ∼12min). Participants reported switches in perception through a button press. We considered that task would involve a relatively lower phasic response over a relatively larger tonic tone (high-tonic LC). **(C)** The auditory salience detection task consisted of an oddball paradigm requiring button-press reports on the perception of rare deviant target tones (20% occurrence) within a stream of frequent tones (total duration ∼10min). We considered that task would involve a relatively larger phasic response over a lower relatively tonic tone (low-tonic LC).

Younger participants completed the fMRI session immediately following the baseline night but older participants were initially part of a different study[19,20] and completed the MRI procedures in addition to their initial engagement, which included the baseline night recording. Consequently, the baseline nights of sleep and MRI sessions were completed about 1.25y apart (mean±SD: 15.5±5.3 months). Prior to the fMRI session, older participants slept regularly for 1 week and were maintained in dim light (<10 lux) for 45 minutes.

### Sleep EEG metrics

Sleep was staged in 30s-epochs using an automatic algorithm (ASEEGA, PHYSIP, Paris)[21] to provide REM sleep percentage and total sleep time (TST). Averaged power was computed per 30-min bin, adjusting for the proportion of rejected data and it was subsequently aggregated in a sum for REM[22] to provide REM theta energy (overnight cumulated 4.25-8Hz power). Sigma power (12.25–16 Hz) prior to REMS was calculated as the mean sigma power before REMS episodes, obtained by dividing the total sigma power in the 1-minute period preceding each REMS episode by the number of REMS episodes. Sigma power was computed as the weighted sum of 4s artifact-free windows (2s overlap per 30s epoch).

### Cognitive tasks

#### Visual perceptual rivalry task – <u>high-tonic LC</u>

The task consisted of watching a 3D Necker cube, which can be perceived in two different orientations (**Figure 1B**), for 10 blocks of 1min separated by 10s of screen-center cross fixation (∼12min). Participants were instructed to report switches between the two percepts through a button press. Given the top-down nature of the task, we considered that if the task recruited the LC, it would trigger a relatively lower phasic response over a relatively larger tonic tone (high-tonic LC).

#### Auditory salience detection task – <u>low-tonic LC</u>

The task consisted of an oddball paradigm requiring reports on the perception of rare deviant target tones (1,000Hz, 100ms, 20% of tones) that were pseudo-randomly interleaved within a stream of standard stimuli (500Hz, 100ms) through a button press (**Figure 1C**). The task included 270 stimuli (54 targets; ∼10min). Given the bottom-up nature of the task, we considered that if the task recruited the LC, it would trigger a relatively larger phasic response over a relatively lower tonic tone (low-tonic LC).

Pupil size was recorded (Eyelink-1000, SR Research, Osgoode, ON, Canada; sampling rate: 1000Hz) during both cognitive tasks only in a subset of participants due to technical difficulty in maintaining stable signal (in part because of the small window between receive and transmit parts of the MR head coils through which the eye is within camera view) leading to missing/corrupted data (> 25% of missing/corrupted). Analyses of pupil data of the perceptual rivalry and salience detection tasks respectively included 32 (23 women; 23 young individuals) and 27 (20 women; 19 young individuals) individuals with 22 participants common to both tasks (15 women; 15 young individuals).

#### MRI data acquisition and preprocessing

MRI data were acquired using a MAGNETOM Terra 7T MRI system (Siemens Healthineers, Erlangen, Germany). FMRI and sMRI data were preprocessed using SPM12, ANTs and SynthStrip brain extraction tool[23], as fully described previously[13]. The preprocessed data were resampled to a 1mm^3^ resolution. Individual statistical analyses included one regressor of interest consisting of a switch in perception (perceptual rivalry) or the occurrence of a target deviant tone (salience detection) modeled as an event (convolved with the canonical hemodynamic response function - HRF). Participant movement, respiration, and heart rate were used as covariates of no interest (physiological data of 4 volunteers were not available).

Individual LC masks were manually delineated by 2 experts based on LC-specific images (as in[13]) to extract LC activity estimate or LC responses to the task (i.e., in arbitrary units – a.u. – as the mean value of the of the multiple regression fits over the LC mask) during both tasks and to compute a LC probabilistic map in the group space for visualization. The T1 structural whole-brain image was used to extract individual total intracranial volume (TIV) using CAT12 toolbox.[24]

### Statistics

Statistical analyses were performed in SAS 9.4 (SAS Institute, NC, USA) and consisted of Generalized Linear Mixed Models (GLMM) with 4 main sleep features of interest (REM sleep latency, REM sleep percentage, REM Theta energy, sigma power prior to each REM sleep episode) as the dependent variables (with adjusted distribution), LC activity as an independent variable, as well as age group (younger and older individuals), sex, TST and TIV. Outliers among all variables lying beyond four times the standard deviation were removed from the analysis (maximum 2 data points were removed). The final number of individuals included in each analysis is reported in **table 2** and **table 3**. Partial R^2^ *(R*^2*^) values were computed as previously described.[13] We used Pearson correlations to assess the association between transient pupil dilation and estimates of LC activity and for visualization of the GLMM outputs. The analyses included 4 main dependent variables of interest: Benjamini and Hochberg False Discovery Rate (FDR) correction considering 4 independent tests was used to test for significant associations [p<.0125 (for rank 1/4); p<.025 (for rank 2/4); p<.0375 (for rank 3/4); p<.05 (for rank 4/4)]. All models included an interaction term between LC activity and age group. If the Bayesian Index Criterion (BIC) of the model without the interaction was better (i.e. lower) and the interaction term was not significant it was removed from the model.

**Table 2.** Associations between REM sleep metrics and LC activity estimated via <u>the visual perceptual</u> <u>rivalry (high-tonic) task</u>.

| Sleep metric<br>(dependent<br>variable) | LC activity | Age group | Sex | TIV | Total sleep time |
| --- | --- | --- | --- | --- | --- |
| **REM Theta**<br>**energy**<br>**(N=51)** | F(1,45)=11.39<br>**P=0.001\***<br>**R<sup>2</sup>=0.201** | F(1,45)=0.14<br>P=0.709 | F(1,45)=0.22<br>P=0.645 | F(1,45)=0.08<br>P=0.781 | F(1,45)=5.15<br>**P=0.028**<br>**R<sup>2</sup>=0.102** |
| **Sigma power**<br>**prior to REM**<br>**(N=52)** | F(1,46)=5.55<br>**P=0.023\***<br>**R<sup>2</sup>=0.107** | F(1,46)=0.60<br>P=0.442 | F(1,46)=3.64<br>P=0.062 | F(1,46)=2.18<br>P=0.147 |  |
| **REMS**<br>**percentage**<br>**(N=52)** | F(1,47)=2.05<br>P=0.159 | F(1,47)=13.27<br>**P=0.0007**<br>**R<sup>2</sup>=0.220** | F(1,47)=1.66<br>P=0.204 | F(1,47)=3.15<br>P=0.082 |  |
| **REMS latency**<br>**(N=52)** | F(1,46)= 0.10<br>P= 0.750 | F(1,46)= 2.01<br>P= 0.163 | F(1,46)= 1.27<br>P= 0.265 | F(1,46)= 0.27<br>P= 0.602 | F(1,46)= 0.74<br>P= 0.392 |

Prior to the analysis, we removed the outliers among all variables by excluding the samples lying beyond four times the standard deviation (the final number of individuals included in each analysis is reported below each dependent variable).
\* Significant following Hochberg False Discovery Rate (FDR) correction for 4 independent tests (see methods)
In all models the interaction between LC activity and age group was not significant. Goodness of fit metric (BIC) indicated that the interaction term should be removed. LC: locus coeruleus; TIV: total intracranial volume; REM: rapid eye movement; REMS: rapid eye movement sleep.

**Table 3.** Associations between REM sleep metrics and LC activity estimated via <u>the auditory</u> <u>salience detection (low-tonic) task</u>.

| Sleep metric<br>(dependent<br>variable) | LC activity | Age group | LC<br>activity*age<br>group | Sex | TIV | Total sleep<br>time |
| --- | --- | --- | --- | --- | --- | --- |
| **REM Theta**<br>**energy**<br>**(N=51)** | F(1,44)=6.74<br>**P=0.012\***<br>**R<sup>2</sup>=0.132** | F(1,44)=0.54<br>P=0.468 | F(1,44)=7.19<br>**P=0.010\***<br>**R<sup>2</sup>=0.140** | F(1,44)=0.14<br>P=0.707 | F(1,44)=0.10<br>P=0.750 | F(1,44)=3.48<br>P=0.068 |
| **Sigma power**<br>**prior to REM**<br>**(N=50)** | F(1,44)=1.90<br>P=0.175 | F(1,44)=0.26<br>P=0.611 | F(1,44)=2.35<br>P=0.132 | F(1,44)=8.01<br>**P=0.007**<br>**R<sup>2</sup>=0.154** | F(1,44)=3.11<br>P=0.084 |  |
| **REMS**<br>**percentage**<br>**(N=52)** | F(1,46)=1.38<br>P=0.245 | F(1,46)=15.60<br>**P=0.0003**<br>**R<sup>2</sup>=0.253** | F(1,46)=2.07<br>P=0.157 | F(1,46)=0.23<br>P=0.635 | F(1,46)=5.56<br>**P=0.022**<br>**R<sup>2</sup>=0.107** |  |
| **REMS latency**<br>**(N=52)** | F(1,45)= 2.88<br>P= 0.096 | F(1,45)= 0.75<br>P= 0.389 | F(1,45)=1.40<br>P=0.242 | F(1,45)= 2.07<br>P= 0.157 | F(1,45)= 0.88<br>P= 0.351 | F(1,45)= 0.96<br>P= 0.331 |

Prior to the analysis, we removed the outliers among all variables by excluding the samples lying beyond four times the standard deviation (the final number of individuals included in each analysis is reported below each dependent variable).
\* Significant following Hochberg False Discovery Rate (FDR) correction for 4 independent tests (see methods)
In all models, the results without the interaction between LC activity and age group were not significant. Goodness of fit metric (BIC) indicated that the interaction term provides a better fitness.
LC: locus coeruleus; TIV: total intracranial volume; REM: rapid eye movement; REMS: rapid eye movement sleep.

Importantly, statistical analyses are slightly modified compared to[13] with the inclusion of TIV as a covariate (rather than BMI) to take into account non-specific variation in EEG power computations. Statistical outcomes related to the auditory salience detection task may therefore slightly vary compared with our previous publication.[13]

We computed a prior sensitivity analysis to get an indication of the minimum detectable effect size in our main analyses given our sample size. According to G*Power 3 (version 3.1.9.4),[25] taking into account a power of .8, an error rate α=.05, and a sample size of 52, we could detect medium effect sizes *r*>.39 (2-sided; CI:.13–.6; *R*²>.15, CI:.02–.36) within a linear multiple-regression framework including 2 tested predictor (LC activity, age group) and 2/3 covariates (sex, TST, TIV where relevant).

## Results

Fifty-two healthy individuals aged 18 to 31y (N=34; 28 women) and 50 to 70y (N=18; 14 women) completed an fMRI protocol consisting of a top-down visual perceptual rivalry task[26], which we considered could trigger a lower phasic response of the LC over a relatively larger tonic tone (high-tonic LC), and a bottom-up auditory salience detection task[27], which we considered could trigger a larger phasic response of the LC over a relatively lower tonic tone (low-tonic LC task). Both tasks successfully triggered a phasic response of the left LC when inspecting the whole-brain statistical analyses over the entire sample (**Figure 2A and B**). The recruitment of the LC was further supported by the fact that, in a subset of participants (see methods), we successfully detected significant transient pupil dilation around the event of interest (**Figure 2C, D, E and G**). The LC is indeed considered to drive part of the transient pupil dilation associated with the detection of salient stimuli or with changes in perception in a perceptual rivalry context.[27,28]

**Figure 2.**
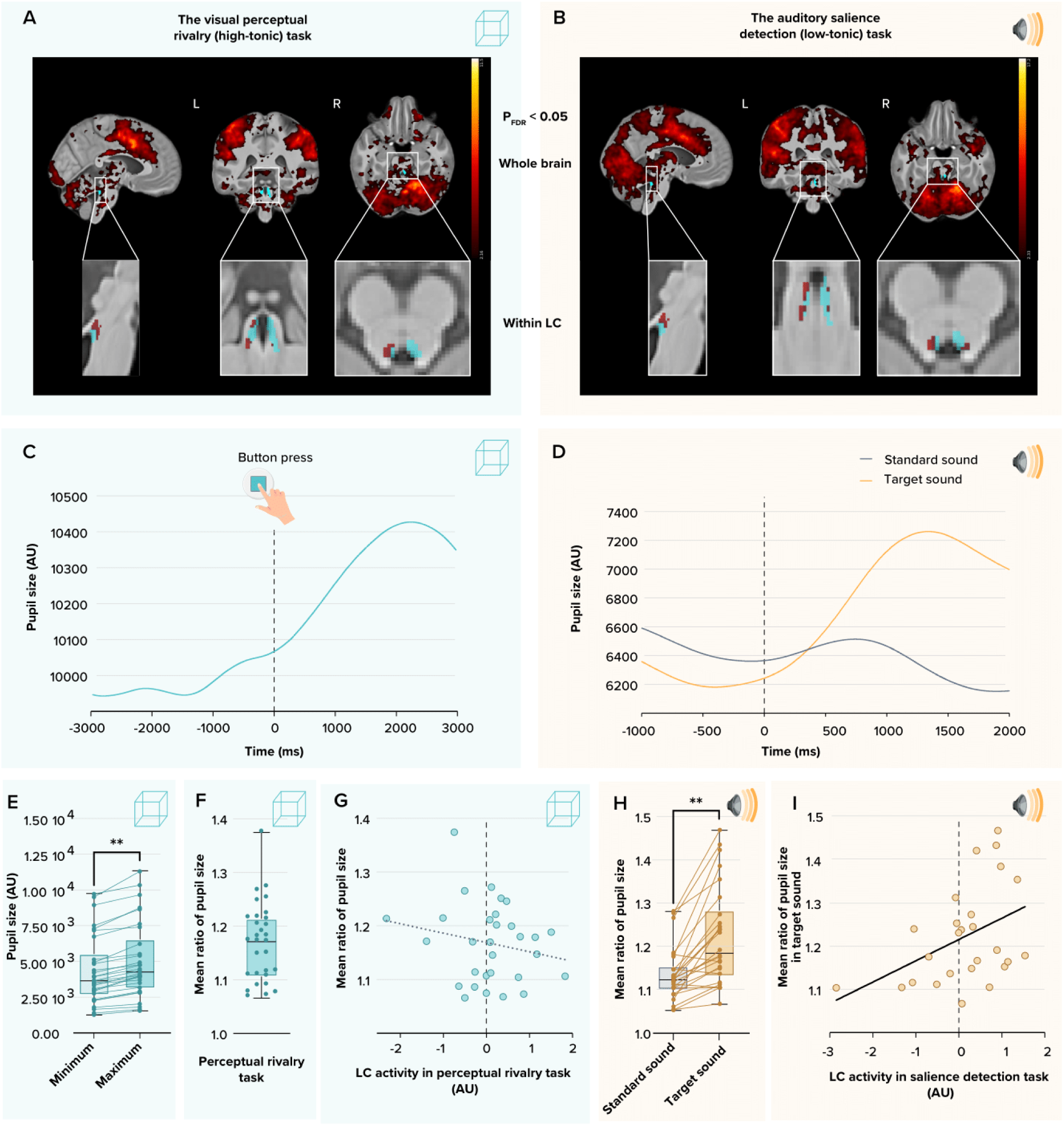
**(A)** Whole-brain and LC responses to the perceptual switches during the visual perceptual rivalry task. Sagittal, coronal, and axial views [MNI coordinates: (−4 −37 -21 mm)]. The images at the top show the whole-brain results using significance for a threshold of p< 0.05 FDR-corrected (t > 2.16) over the group average brain structural image coregistered to the MNI space. Insets at the bottom show the LC probabilistic template (blue) created based on individual LC masks and the significant activation detected within this mask (dark grey). **(B)** Whole-brain and LC responses to the target sound during the auditory salience detection task. Sagittal, coronal, and axial views [MNI coordinates: (−4 −34 -21 mm)]. The images at the top show the whole-brain results using significance for a threshold of p< 0.05 FDR-corrected (t > 2.33) over the group average brain structural image coregistered to the MNI space. Insets at the bottom show the LC probabilistic template (blue) and the significant activation detected within this mask (dark grey). The legend shows the t-values associated with color maps. **(C)** A representative example of the pupil dilation in response to the perceptual switches during the visual perceptual rivalry task in one participant. **(D)** A representative example of the pupil dilation in response to the standard and target sounds during the auditory salience detection task in the same subject as in panel C. **(E)** Minimum and maximum pupil size before and after the perceptual switches during the visual perceptual rivalry task. Maximum pupil size was significantly higher than the minimum (N=32, p < 0.0001). Arbitrary units (AU) of the pupil size in the figures C, D and E are based on pixel number following proprietary algorithms application yet do not correspond exactly to pixel number. **(F)** Mean ratio of the pupil size in the perceptual rivalry task. The mean ratio of pupil size was computed as the change in the pupil diameter from before to after the perceptual switches. **(G)** Association between LC activity during visual perceptual rivalry task and mean ratio of pupil size in response to the perceptual switches. The association was not significant (Pearson’s correlation r=-0.201, P=0.29). **(H)** Mean ratio of pupil size in response to standard sound and target sound during the auditory salience detection task. The mean ratio of pupil size was computed as the change in the pupil diameter from before to after the auditory stimulus presentation. The change in pupil size was significantly higher for target versus standard sound (N=27, p < 0.0001). (**I**) Association between LC activity during auditory salience detection task and mean ratio of pupil size in response to the target sound. We found a significant positive Pearson’s correlation (r=0.4, P=0.03).

We therefore extracted the activity of the left LC in all individuals. We first found that transient pupil dilation during the salience detection task was significantly correlated to the activity estimate of the left LC (Pearson’s r=.4, **p=.03**; **Figure 2H**), while it was not for the perceptual rivalry task (**Figure 2F**, Pearson’s r=-.20, p=.29). Since pupil responses are associated with phasic responses of LC,[29] the absence of significant correlation in the perceptual rivalry task is in line with our hypothesis that this task involved more tonic LC activity. We further found that LC responses during both tasks were not correlated (Figure 3A) (Pearson’s r= .08, p=.54), indicating that the activity of LC in one task did not linearly predict its activity in the other task and supporting that both tasks probe different aspects of LC activity. We did find a significant positive Pearson correlation (whole sample: r=0.357; p=0.010; in the young only: r=0.415; p=0.018; in the older only: r=-0.457, p=0.056) between LC activity estimate during the salience detection task and squared LC activity estimate in the perceptual rivalry task. This suggests a U- shape relationship between LC activity estimate in two tasks and may support the idea that tonic and phasic activities interact in a nonlinear manner. This would deserve future investigations.

**Figure 3.**
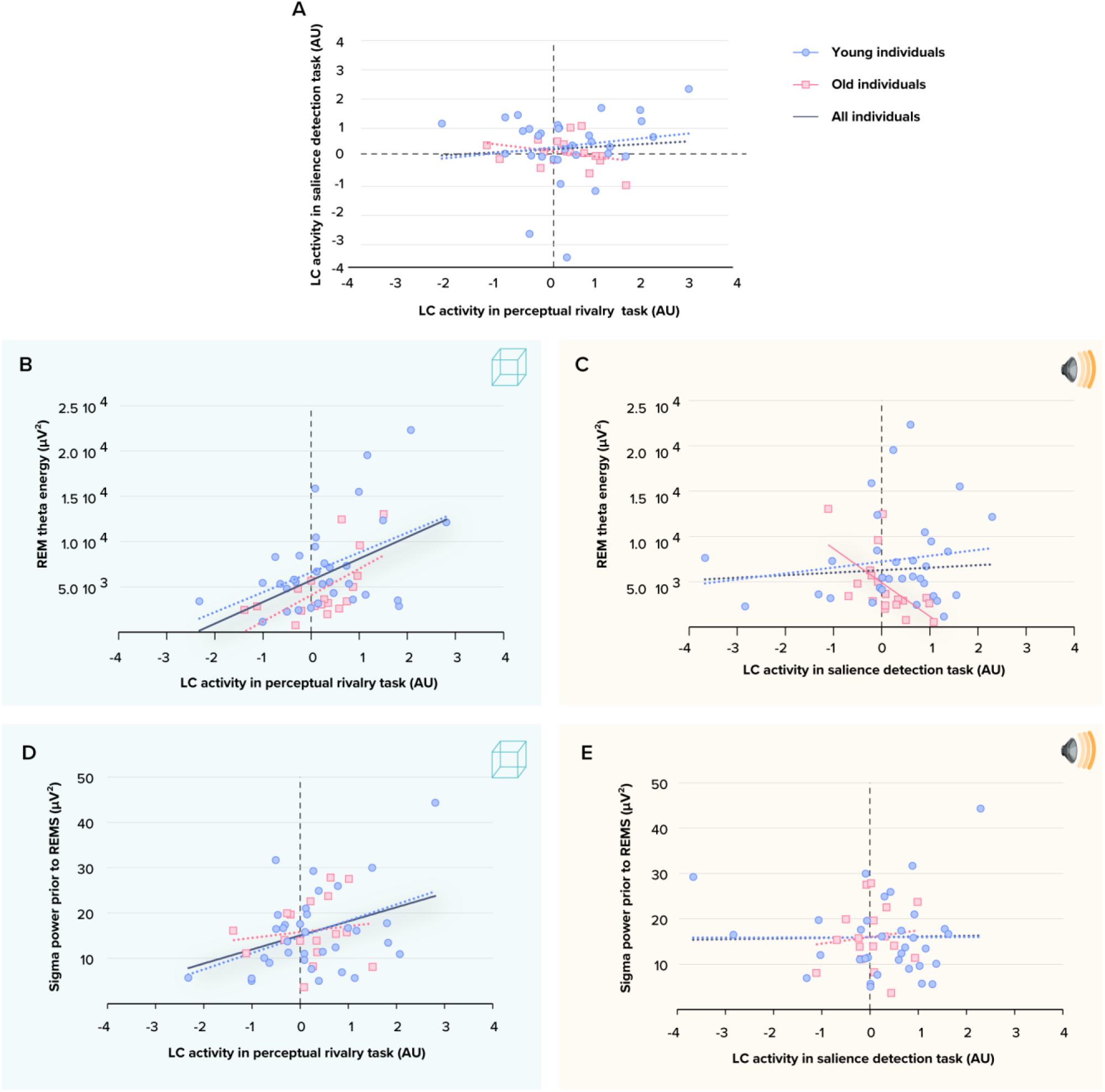
**(A)** LC activity estimates during the visual perceptual rivalry task and auditory salience detection task were not significantly correlated (p=0.54). **(B)** Association between REM theta energy and the LC activity estimates during the visual perceptual rivalry task. The GLMM yielded a significant main effect of LC activity (p=0.0015). **(C)** Association between REM theta energy and the LC activity estimates during the auditory salience detection task. The GLMM yielded a significant age group by LC activity interaction (p=0.0103), and post hoc analyses led to a significant association for the older (p=0.008) but not the young group (p=0.75). **(D)** Association between sigma power prior to REM sleep and the LC activity estimates during the visual perceptual rivalry task. The GLMM yielded a significant main effect of LC activity (p=0.026). **(E)** Association between sigma power prior to REM sleep and the LC activity estimates during the auditory salience detection task. The GLMM showed neither a significant main effect of LC activity (p=0.859) nor a significant age group by LC activity interaction (p=0.132). Simple regression lines are used for a visual display and do not substitute the GLMM outputs. The black line represents the regression irrespective of age groups (young + old, n = 52). Solid and dashed regression lines represent significant and non-significant outputs of the GLMM, respectively.

We then targeted our main goal: testing relationships between LC activity and canonical characteristics of REM sleep estimated based on a night of sleep in the lab under EEG. Considering first the perceptual rivalry task, we found a significant positive association between REM theta energy and high-tonic LC activity (t=3.38; **p= .001**; **Figure 3B**) on top of the main effect of TST, while the other covariates were not significant (**Table 2)**. As previously reported,[13] we then found a significant association between REM theta energy and the interaction between low-tonic LC activity during the salience detection task and age group (t=2.68; **p=.01; Table 3**) that came on top of a main effect of LC activity (t=-2.78; **p=.008**) (**Figure 3C**) while the other covariates were not significant (**Table 3**). Post hoc contrasts revealed that higher REM theta energy was associated with a lower response of the LC in older (t= -2.78; **p=.008**) but not in younger (t=.33; p=.74) individuals, indicating that the main effect of LCactivity was mainly driven by the older participants.

For both low-tonic and high-tonic LC activity the inclusion of REM sleep duration and/or REM sleep percentage instead of or on top of TST did not modify the statistical outputs of the models (data not shown). Furthermore, REM theta energy associations with the activity of the LC during both tasks remain similar when including them simultaneously in the same GLMM, i.e. a significant main effect of high-tonic LC activity (F_(1,41)_=9.52, **p=.0036**) and a weak trend for interaction between low-tonic LC activity and age: F_(1,41)_=2.72, p=.1), supporting that they explain different parts of the variance in REM theta energy (using squared values of LC activity estimates of each task separately or together estimates did not yield significant outputs – not shown).

No association was found when considering the other macroscopic REM sleep metrics (REM sleep onset latency, REM sleep percentage) (**Suppl. Figure S1, Table 2 and 3**). Further analyses indicated that the associations with REM theta energy were specific to this particular band as no significant association was uncovered between the energy of the other spectral bands of the EEG during REM sleep (except for a significant main effect of low tonic LC activity [t= -2.33; p=.025) and of an age-group-by-low tonic LC activity interaction (t= 2.14; p=.037) when using REM alpha energy as the dependent variable - **Suppl. Figure S2, Suppl. Table S1 and S2**]. The significant associations we detected were also specific to REM sleep as NREM slow wave energy (SWE), i.e. the cumulated overnight power over the delta band (.5-4Hz) typical of NREM sleep, was not significantly associated with either low-tonic and high-tonic LC activity (**Suppl. Figure S2, Suppl. Table S1 and S2**).

In the next steps, we turned towards sigma power prior to REM sleep as it has been causally related to LC activity during sleep in animal models.[16] Statistical analyses yielded a significant positive main effect of high-tonic LC activity during the perceptual rivalry task (t=2.36; **p=0.022**; **Table 2**; **Figure 3D**) and no significant interaction between LC activity and age group. In contrast, no significant association with low-tonic LC activity during the salience detection task (p=0.17) and no significant interaction between LC activity and age group were detected (**Table 3**; **Figure 3E**). In both analyses, we did not detect significant effects of the other covariates except for main effect of sex: a statistical trend in the perceptual rivalry task (t=-1.91; p= 0.062; Table 2) and a significant main effect in the salience detection task (t=-2.83; p= 0.007; Table 3), resulting from men presented less sigma power compared to women (Suppl. Figure S4). As it is not the main focus of our study, sex difference will not be discussed in detail. We note that the sex difference in sigma power prior to REM may be related to the reduced percentage of REM sleep sometimes found in other studies [30] (though REM sleep percentage did not differ between sexes in our sample: t = -0.63; p = 0.52).

Finally, when including the activity of both tasks in the same GLMM with sigma power prior to REM as the dependent variable, the analysis yielded a significant main effect of high-tonic LC activity (F_(1,42)_=7.62,**p=.0085**) as well as a trending interaction between low-tonic LC activity and age (F_(1,42)_=3.77, p=.058). As for REM theta energy, this result suggests that both types of LC activity estimates explain different parts of the variance in sigma power prior to REM.

For completeness, the final steps of our analyses consisted of exploratory analyses for associations between the activity of LC estimated during both tasks and other REM sleep metrics, i.e. REM sleep duration, REM bouts duration, and number of arousals during REM sleep. These led to no significant association (**Suppl. Figure S3, Suppl. Table S1 and S2**).

## Discussion

Whether the variability in the functioning of human LC may contribute to the interindividual variability in sleep is not established. In the present cross-sectional study, we show that the activity of the LC, assessed during wakefulness through 2 distinct cognitive tasks, is related to the variability of REM sleep intensity, as reflected by the energy of the theta band of the EEG. We find that the associations depend on the task considered - presumably because they do not rely on the same balance between phasic and tonic LC activity, and on age - when considering LC activity during the low-tonic task. In addition, we find that the amount of sigma band oscillations immediately preceding REM sleep is associated with the activity of the LC assessed during the high-tonic task. These findings add to the existing view that LC function is essential for sleep regulation that was mainly brought by animal experiments. We show that the *in vivo* variability in the activity of the human LC may govern part of the variability in sleep quality assessed using electrophysiology. The findings may have implications for the neuropathology of several brain disorders such as insomnia, Alzheimer’s disease (AD), and Parkinson’s disease (PD) for which the LC may play a pivotal role through sleep.

LC cells discharge action potentials in both tonic and phasic modes during wakefulness. The tonic mode consists of irregular but constant baseline activity (1-6 spikes/s), while phasic activity is characterized by short (<300ms) bursts of high-frequency activity followed by a long period of sustained inhibition of spontaneous activity.[31] A balanced tonic activity is required for an optimal arousal level allowing the expression of phasic activity in response to salient stimuli or behavioral changes.[15] While the LC was considered for long as a quiet region during sleep, recent animal studies revealed that the activity of the LC varies importantly during NREM sleep and may even transiently reach activity levels similar to wakefulness.[10] NE level, driven by LC neuron activity, fluctuates during NREM sleep between periods of higher and lower NE tone following an infraslow rhythm of ∼50s.[10] This periodic decrease in NE tone must reach levels that are low enough to open the gate for REM sleep, which is considered to require sustained NE-free periods to allow, for instance, synaptic pruning.[4] The periodic decrease of the LC activity is reflected in periodic increases in the expression of sleep spindles and in sigma power such that each REM episode is preceded by a pronounced transient increase in sigma power[32] that is well known to somnologists. The overall picture emerging from this recent literature is that LC activity shapes the macro- and microstructure of sleep and that inappropriate fluctuations in LC tonic activity can disturb sleep and in particular REM sleep.[4,10]

We find that a higher phasic response of the LC during the top-down perceptual rivalry task, arguably characterized by a relatively high tonic activity of the LC is associated with a larger expression of theta oscillations during REM sleep and of sigma power prior to REM sleep episodes. We interpret this finding as the reflection of a more appropriate lower tonic activity of the LC which allows a better expression of its phasic response during the task. In contrast, if tonic activity during the task is too high, it partly masks the phasic response of the LC. We detect this during wakefulness but surmise that it constitutes a trait such that the more appropriate tonic activity of the LC present during wakefulness would also be present during sleep. This would mean that, in the case of a low tonic LC activity allowing for a large dynamic range of phasic LC oscillation, NE tone reaches a lower level prior to a REM sleep episode, allowing a large expression of sigma oscillation and the initiation of a REM sleep episode and complete silencing during REM. This would in turn enhance the expression (intensity) of REM theta energy, the most typical oscillatory mode of REM sleep. Theta activity during REM sleep is considered to reflect hippocampus activity and plays key roles in memory consolidation during REM sleep.[33] The putative more appropriate tonic activity of the LC we report could therefore reflect a better expression of REM sleep functions.

In contrast, we find that a larger expression of the phasic response of the LC during the bottom-up salience detection task, which is arguably underlined by a lower tonic activity of the LC, is associated with lower expression of theta oscillations during REM sleep, particularly in older individuals. We link our results to the recent report in rodents that the so-called startle effect, which is triggered by abrupt sensory changes, is attenuated under higher vs. lower arousal levels as well as when LC activity is induced prior to the salient stimulation.[34] This was in line with the established observation that elevated tonic firing of the LC-NE neurons makes the LC sensory-evoked response less pronounced.[31] This would mean that in individuals with lower tonic activity of the LC during wakefulness, the dynamic range of the phasic response of the LC is potentially large. This trait-like phasic LC activity could become maladaptive and during sleep, perturb the activity of limbic areas, thereby resulting in a reduction in REM theta oscillations.

This process becomes particularly manifest in aging which is characterized by increased sleep fragility and degradation of sleep quality.[35] Overall, we interpret our findings as a reflection of the requirement for a balanced tonic LC activity for optimal REM sleep. This notion is supported, in our view, by the fact that our statistical outputs remain similar (either significant or as weak trends) when including concomitantly the activity of the LC as assessed in a low-tonic and high-tonic context. Our findings may further be related to the diversity in neuron types within the LC that is emerging in the literature, for instance, if both tasks were recruiting different proportions of LC neurons co- expressing or not dopamine or projecting to different brain territories[6] and were therefore characterizing the activity of these different neurons during sleep.

We stress that, although plausible, our interpretations remain putative and that measures of LC activity during sleep are required to confirm our hypotheses. Future studies could assess the connectivity of the LC with other brain structures in humans, as sleep and wakefulness regulation relies on a network of subcortical nuclei rather than on the LC alone.[36] Furthermore, while we interpret our finding in “LC centric” manner, i.e. the activity of the LC during wakefulness reflects its activity during sleep, it may also be that the activity of the LC during wakefulness shapes the activity of other brain areas during sleep. Sleep oscillations depend indeed in prior wakefulness in a regional use-dependent manner[37] such that the impact of NE tone and LC activity on other regions, and particularly over cortical regions, may be reflected in a larger or lower expression of theta oscillation during REM.

Beyond the healthy population, our findings are in line with the view that inappropriate LC activity during both wakefulness and sleep can contribute to insomnia disorder.[4] Insomnia is characterized by a hyperarousal state that is presumably due in part to inappropriately high tonic activity of the LC during wakefulness, contributing to delayed sleep onset, and during sleep, contributing to early and/or multiple awakenings. This would also lead to disturbed REM sleep and unresolved emotional distress.[4] Our participants being healthy and devoid of sleep disorders, our finding could reflect the continuum between normal sleep and insomnia. They could also be indicative of the neuropathological mechanisms eventually leading to insomnia, i.e. a prolonged inappropriate LC tonic activity and REM sleep disturbance are consequences or contribute to the progressive changes leading to insomnia or maintaining the disorder. LC is also central to several neurodegenerative diseases characterized by preclinical alteration in sleep. The LC is indeed among the first sites showing abnormal tau and alpha-synuclein aggregates, hallmarks of AD and PD, respectively.[38,39] Abnormal tau inclusions can even be detected post-mortem as early as during the second decade of life[38] and were suggested to be associated with larger cortical excitability in late middle-aged healthy individuals.[40] In addition, *post-mortem* degeneration of the human LC was recently associated with alteration in the rest-activity cycle over the ∼7y preceding death, particularly in those with higher cortical AD pathology.[41] Future research could try to link our findings to assessments of tau and alpha-synuclein burden within the LC, though the level of protein aggregate would go undetected *in vivo* in young individuals, and would remain challenging to assess over such a deep and small nuclei in older individuals.

As previously mentioned,[13] although our study provides new insights into the associations between LC activity and sleep variability in humans, it bears limitations. The baseline night of sleep was followed by the fMRI acquisition the next day in young individuals, while in the older group, there was a ∼1y gap between the baseline night and the fMRI session. Although sleep changes over the lifespan,[42] it is stable over a short life period (e.g. a few years).[43] We therefore consider that this important limitation is unlikely to fully explain our results, particularly given the fact that no age-related differences were found for the high-tonic perceptual rivalry task. Moreover, our sample was primarily composed of women. Although it was considered in our statistical analyses, this limits the generalizability of our results. In addition, although it represents a considerable data collection effort, the size of our sample remains modest, particularly for the older subsample. Replication in a larger sample is therefore warranted. Future studies could also apply individually tailored HRF to assess LC response. We used the canonical HRF to model activity over the entire brain to model average LC response, while individual LC responses can differ across individuals.[44] Likewise, we used multiple linear regression approaches for the statistical, mostly precluding isolation of non-linear relationships. Finally, the LC activity was measured using task-evoked fMRI BOLD signals, which reflect changes in neural activity relative to a baseline condition. Although, on average responses were positive (cf. Figure 2 A-B), the negative values in LC activity detected in some participants (cf. Figure 2 G & I) represent instances where the BOLD signal in the LC during a task condition is lower than the baseline signal. The meaning of negative BOLD value is unclear. They may reflect differences in individual task engagement, arousal levels, or baseline LC activity. Further investigation is needed to better understand the underlying physiological or methodological factors contributing to this phenomenon.

## Conclusions

In summary, we provide original evidence that, aside from its integrity, the functioning of the LC shapes part of the quality of sleep. Our data suggest that an optimal level of LC tonic activity is required for an optimal expression of REM sleep and its functions. This reinforces the view that the LC-NE is promising for interventions aiming at improving sleep quality, including preventing and/or delaying brain disorders. Our findings suggest that these interventions could aim at restoring optimal LC functioning during sleep as well as during wakefulness.

## Supporting information

SUPPLEMENTARY MAERIAL

## List of abbreviations

AD: Alzheimer’s disease
BAI: Beck Anxiety Inventory
BDI: Beck Depression Inventory
BMI: Body Mass Index
CI: confidence intervals
EEG: electroencephalogram
ESS: Epworth Sleepiness Scale
FDR: False Discovery Rate
fMRI: functional Magnetic Resonance Imaging
GLMM: Generalized Linear Mixed Models
LC: locus coeruleus
NE: norepinephrine
PD: Parkinson’s disease
PSQI: Pittsburgh Sleep Quality Index
REM: Rapid Eye Movement
REMS: rapid eye movement sleep.
SD: standard deviation
sMRI: structural MRI
SWE: slow wave energy
TIV: Total Intracranial Volume
TST: Total Sleep Time

## Declarations

### Ethics approval and consent to participate

This study was approved by the faculty-hospital ethics committee of ULiège. All participants provided written informed consent and received financial compensation.

### Consent for publication

Not applicable.

### Availability of data and materials

The processed data and analysis scripts supporting the results included in this manuscript are publicly available via the following open repository: https://gitlab.uliege.be/CyclotronResearchCentre/Public/fasst/Balanced_LC_activity_and_REM_sleep. The raw data could be identified and linked to a single subject and represent a large amount of data. Researchers willing to access to the raw should send a request to the corresponding author (GV). Data sharing will require evaluation of the request by the local Research Ethics Board and the signature of a data transfer agreement (DTA).

### Authors’ contributions

Study concept and design by NM, GV and PM. Data acquisition and analysis by NM, PT, EK, RS, EB, AB, IC, IP, FB, ID & SS. Methodological support and/or support in the interpreting the data by CBe, CBa, CP, PM, FC, MZ, LL. Funding was mostly obtained by CBa, FC, CP, PM and GV. NM and GV drafted the first version of the manuscript. All authors revised the manuscript and had final responsibility for the decision to submit for publication.

## Acknowledgments

This work was conducted at the GIGA-In Vivo Imaging platform of ULiège, Belgium. We thank N. Beliy, E. Lambot, C. Hagelstein, S. Laloux, A. Claes, C. Degueldre, B. Herbillon, G. Hammad, P. Hawotte, B. Lauricella, T. Pontus, T. Gendron, J. Read, A. Robert, H.I.L. Jacobs, A. Lesoinne, A. Deward and Illumine company for their help in the different stages of the study.

