## SUPPLEMENTARY MAERIAL for "REM sleep quality is associated with balanced tonic activity of the locus coeruleus during wakefulness"

### Supplementary Methods

*Participant*

Due to a miscalculation at screening, 1 older participant had a BMI of 30.9 and one of the younger participants had a BMI of 28.4. Since their data do not deviate substantially from the rest of the sample these participants were included in the analyses (including BMI as a covariate in our statistical models did not modify our results).

*Protocol*

The evening before the baseline night, participants arrived at the laboratory 3 hours before their habitual bedtime, completed questionnaires including Beck Depression Inventory (BDI)[1], Beck Anxiety Inventory (BAI)[2], the Pittsburgh Sleep Quality Index (PSQI)[3], and Epworth sleepiness scale (ESS)[4] for assessing depression, anxiety, sleep quality, and sleepiness, respectively.

The procedures for the baseline night recordings were identical in older and younger participants. Prior to the fMRI session, older participants slept regularly for 1 week (verified with a sleep diary; based on our experience, actigraphy reports and sleep diaries do not deviate substantially in older individuals). Older participants were maintained in dim light (<10 lux) for 45min before the fMRI scanning.

*Sleep EEG metrics*

Eleven channels were used for the baseline night (F3,z,4; C3,z,4; P3,z,4; O1,2) initially referenced to the left mastoid prior to rereferencing offline to the average of both mastoids. Arousals and artefacts were detected automatically[5] to provide the number of arousals during REM sleep, and excluded from the power spectral density analyses. Only frontal electrodes were considered because the frontal region is most sensitive to sleep pressure manipulations;[6] focusing on the frontal electrodes may also facilitate interpretation of future large-scale studies using ambulatory EEG, often restricted to frontal electrodes.

*Cognitive tasks*

*Auditory salience detection task – low-tonic LC.* The recording started with a setting of the volume to ensure an optimal auditory perception.

*MRI data acquisitions and preprocessing*

MRI data were acquired using a MAGNETOM Terra 7T MRI system (Siemens Healthineers), with a single-channel transmit and 32-receiving channel head coil (1TX/32RX, Nova Medical, Forchheim, Germany). Blood-oxygen-level-dependent (BOLD) fMRI data were acquired using a multi-band gradient-recalled echo–echo-planar imaging (GRE-EPI) sequence (main parameters: repetition time=2.340ms, flip angle=90°, matrix size = 160 × 160, 86 axial 1.4 mm–thick slices, MB acceleration factor = 2, GeneRalized Autocalibrating Partial Parallel Acquisition (GRAPPA) acceleration factor = 3, voxel size = 1.4 × 1.4 × 1.4 $mm^3$). The cardiac pulse and the respiratory movements were recorded concomitantly using, respectively, a pulse oximeter and a breathing belt (Siemens Healthineers). The fMRI acquisition was followed by a 2D GRE field mapping sequence to assess B0 magnetic field inhomogeneities with the following parameters: TR = 5.2 ms, TEs = 2.26 ms and 3.28 ms, flip angle (FA) = 15°, bandwidth = 737 Hz/pixel, matrix size = 96 × 128, 96 axial slices, voxel size = 2 × 2 × 2 $mm^3$, acquisition time = 1:38 minutes.

A Magnetization-Prepared with 2 RApid Gradient Echoes (MP2RAGE) sequence was used to acquire T1 anatomical images: TR = 4,300 ms, TE = 1.98 ms, FA = 5°/6°, TI = 940 ms/2,830 ms, bandwidth = 240 Hz/pixel, matrix size = 256 × 256, 224 axial 0.75 mm–thick slices, GRAPPA acceleration factor = 3, voxel size = 0.75 × 0.75 × 0.75 $mm^3$, acquisition time = 9:03 minutes.[7] The LC-specific sequence consisted of a 3D high-resolution magnetization transfer–weighted turbo-flash (MT-TFL) sequence with the following parameters: TR = 400 ms, TE = 2.55 ms, FA = 8°, bandwidth = 300 Hz/pixel, matrix size = 480 × 480, number of averages = 2, turbo factor = 54, magnetization transfer contrast (MTC) pulses = 20, MTC FA = 260°, MTC RF duration = 10,000 µs, MTC inter-RF delay = 4,000 µs, MTC offset = 2,000 Hz, voxel size = (.4 × .4 × .5)$mm^3$, acquisition time = 8:13 minutes. Sixty axial slices were acquired and centered for the acquisitions perpendicularly to the rhomboid fossa (i.e., the floor of the fourth ventricle located on the dorsal surface of the pons).[8]

The LC appearing hyperintense on the MT-TFL images was manually delineated by 2 expert raters (as in[9]), and the intersection of their masks was computed as the final LC mask for each individual. The masks were used for extracting the LC activity during each task in the participant brain space using the REX Toolbox (https://web.mit.edu/swg/software.htm) (average over all mask voxels per hemisphere) and to compute a LC probabilistic map in the MNI space for group level visualization.

Visualization of whole-brain results over the entire sample was completed following normalization of fMRI and sMRI data to the Montreal Neurological Institute (MNI) space. Due to the small size of the nucleus, LC activation was not expected to survive stringent whole-brain family-wise error (FWE) correction for multiple comparisons. Therefore, a false discovery rate (FDR) correction was conducted using SPM12 to detect voxel-level $p<.05$ results within the LC mask.

*Eye tracking data*

Pupil data were processed as described in[10]. Transient pupil dilation in response to target and standard stimuli (salience detection) consisted of the maximum value detected over the 1.5s following stimulus onset relative to the min pupil diameter over -1000ms to -50ms window preceding stimuli onset. Individual values consisted of the mean of these dilation values per stimulus type. Because of the likely intra- and inter-individual variability in the temporal association between the motor response and the perceptual switch, transient pupil dilation associated with perceptual switches in the perceptual rivalry task was computed differently. Baseline pre-switch pupil size consisted of the median value over a 500ms window centered around the minimum pupil value detected over a window of 3000 ms preceding each button press. The pupil size associated to the switch consisted of the median value over a 500ms window centered around the maximum pupil size value detected over the 3000ms following the baseline minimum pupil values. Individual values consisted of the mean of these trial values. Pupil size associated with the switch were compared to baseline pre-switch pupil size to test for pupil dilation.

**Supplementary Table S1. Exploratory analysis on the associations between sleep metrics and LC activity estimated via <u>the visual perceptual rivalry (high-tonic) task</u>.**

| Sleep metric (dependent variable) | LC activity | Age group | Sex | TIV | Total sleep time |
|---|---|---|---|---|---|
| **REM Delta energy (N=52)** | F(1,46)=3.81 P=0.057 | F(1,46)=0.14 P=0.713 | F(1,46)=0.55 P=0.461 | F(1,46)=0.01 P=0.940 | F(1,46)=0.69 P=0.410 |
| **REM Sigma energy (N=52)** | F(1,46)=1.73 P=0.194 | F(1,46)=1.27 P=0.265 | F(1,46)=0.59 P=0.445 | F(1,46)=0.46 P=0.502 | F(1,46)=5.67 **P=0.021** **$R^{2*}$=0.109** |
| **REM Alpha energy (N=51)** | F(1,45)=3.06 P=0.087 | F(1,45)=0.32 P=0.576 | F(1,45)=0.00 P=0.944 | F(1,45)=1.04 P=0.313 | F(1,45)=6.80 **P=0.012** **$R^{2*}$=0.131** |
| **REM Beta energy (N=51)** | F(1,45)=0.28 P=0.596 | F(1,45)=0.04 P=0.850 | F(1,45)=3.64 P=0.062 | F(1,45)=3.04 P=0.088 | F(1,45)=9.87 **P=0.003** **$R^{2*}$=0.179** |
| **NREM SWE (N=52)** | F(1,46)=2.83 P=0.099 | F(1,46)=0.00 P=0.9634 | F(1,46)=0.02 P=0.882 | F(1,46)=0.35 P=0.556 | F(1,46)=0.14 P=0.709 |
| **REMS duration (N=52)** | F(1,46)=1.56 P=0.217 | F(1,46)=4.76 **P=0.034** **$R^{2*}$=0.093** | F(1,46)=1.48 P=0.229 | F(1,46)=3.70 P=0.060 | F(1,46)=19.85 **P<0.0001** **$R^{2*}$=0.301** |
| **REM bouts duration (N=52)** | F(1,46)=0.51 P=0.478 | F(1,46)=13.24 **P=0.0007** **$R^{2*}$=0.223** | F(1,46)=1.43 P=0.237 | F(1,46)=0.94 P=0.336 | F(1,46)=0.17 P=0. 0.686 |
| **REM bouts number (N=52)** | F(1,46)=0.11 P=0.736 | F(1,46)=3.27 P=0.076 | F(1,46)=0.04 P=0.844 | F(1,46)=0.01 P=0.915 | F(1,46)=6.65 **P=0.013** **$R^{2*}$=0.126** |
| **Number of arousals in REMS (N=52)** | F(1,46)=0.60 P=0.443 | F(1,46)=0.38 P=0.539 | F(1,46)=5.55 **P=0.022** **$R^{2*}$=0.107** | F(1,46)=4.31 **P=0.043** **$R^{2*}$=0.085** | F(1,46)=1.82 P=0.184 |

Prior to the analysis, we removed the outliers among all variables by excluding the samples lying beyond four times the standard deviation (the final number of individuals included in each analysis is reported below each dependent variable).

In all models the interaction between left LC activity and age group was not significant. Goodness of fit metric (BIC) indicated that the interaction term should be removed.

LC: locus coeruleus; TIV: total intracranial volume; REM: rapid eye movement; NREM: Non-rapid eye movement; SWE: slow wave energy; REMS: rapid eye movement sleep.

**Supplementary Table S2. Exploratory analysis on the associations between REM sleep metrics and LC activity estimated via <u>the auditory salience detection (low-tonic) task</u>.**

| **Sleep metric (dependent variable)** | **LC activity** | **Age group** | **LC activity*age** group | **Sex** | **TIV** | **Total sleep time** |
|---|---|---|---|---|---|---|
| **REM Delta energy (N=52)** | F(1,45)=1.35 P=0.251 | F(1,45)=0.29 P=0.592 | F(1,45)=1.79 P=0.187 | F(1,45)=0.00 P=0.945 | F(1,45)=0.16 P=0.688 | F(1,45)=1.01 P=0.320 |
| **REM Sigma energy (N=52)** | F(1,45)=0.32 P=0.572 | F(1,45)=1.87 P=0.177 | F(1,45)=0.54 P=0.466 | F(1,45)=0.13 P=0.722 | F(1,45)=0.39 P=0.536 | F(1,45)=5.8 **P=0.020** **$R^{2*}$=0.114** |
| **REM Alpha energy (N=51)** | F(1,44)=5.35 **P=0.025** **$R^{2*}$=0.108** | F(1,44)=0.01 P=0.925 | F(1,44)=4.59 **P=0.037** **$R^{2*}$=0.094** | F(1,44)=0.72 P=0.400 | F(1,44)=0.91 P=0.346 | F(1,44)=4.97 **P=0.031** **$R^{2*}$=0.101** |
| **REM Beta energy (N=51)** | F(1,44)=1.46 P=0.233 | F(1,44)=0.04 P=0.849 | F(1,44)=0.02 P=0.881 | F(1,44)=3.36 P=0.073 | F(1,44)=4.06 **P=0.050** **$R^{2*}$=0.084** | F(1,44)=11.39 **P=0.001** **$R^{2*}$=0.205** |
| **NREM SWE (N=52)** | F(1,45)=0.65 P=0.424 | F(1,45)=0.00 P=0.954 | F(1,45)=0.80 P=0.377 | F(1,45)=0.05 P=0.827 | F(1,45)=0.20 P=0.658 | F(1,45)=0.19 P=0.664 |
| **REMS duration (N=52)** | F(1,45)=0.29 P=0.594 | F(1,45)=5.69 **P=0.021** **$R^{2*}$=0.112** | F(1,45)=0.50 P=0.482 | F(1,45)=0.36 P=0.550 | F(1,45)=6.09 **P=0.017** **$R^{2*}$=0.119** | F(1,45)=17.29 **P=0.0001** **$R^{2*}$=0.277** |
| **REM bouts duration (N=52)** | F(1,45)=1.40 P=0.243 | F(1,45)=11.04 **P=0.001** **$R^{2*}$=0.197** | F(1,45)=0.19 P=0.666 | F(1,45)=0.84 P=0.364 | F(1,45)=0.97 P=0.331 | F(1,45)=0.00 P=0.981 |
| **REM bouts number (N=52)** | F(1,45)=3.11 P=0.084 | F(1,45)=1.80 P=0.186 | F(1,45)=1.75 P=0.192 | F(1,45)=0.23 P=0.632 | F(1,45)=0.01 P=0.943 | F(1,45)=5.37 **P=0.025** **$R^{2*}$=0.106** |
| **Number of arousals in REMS (N=52)** | F(1,45)=0.85 P=0.360 | F(1,45)=0.32 P=0.573 | F(1,45)=2.24 P=0.141 | F(1,45)=2.18 P=0.146 | F(1,45)=4.45 **P=0.040** **$R^{2*}$=0.089** | F(1,45)=2.16 P=0.148 |

Prior to the analysis, we removed the outliers among all variables by excluding the samples lying beyond four times the standard deviation (the final number of individuals included in each analysis is reported below each dependent variable).

In all models, the results without the interaction between LC activity and age group were not significant. Goodness of fit metric (BIC) indicated that the interaction term provides a better fitness.

The significant main effect of REM alpha energy may consist of prolongation of the effect detected in the theta band. Since the associations with the other bands were not significant, exploratory specificity analyses suggest that the association is specific to theta oscillation and potentially the surrounding alpha band.

LC: locus coeruleus; TIV: total intracranial volume; REM: rapid eye movement; NREM: Non-rapid eye movement; SWE: slow wave energy; REMS: rapid eye movement sleep.

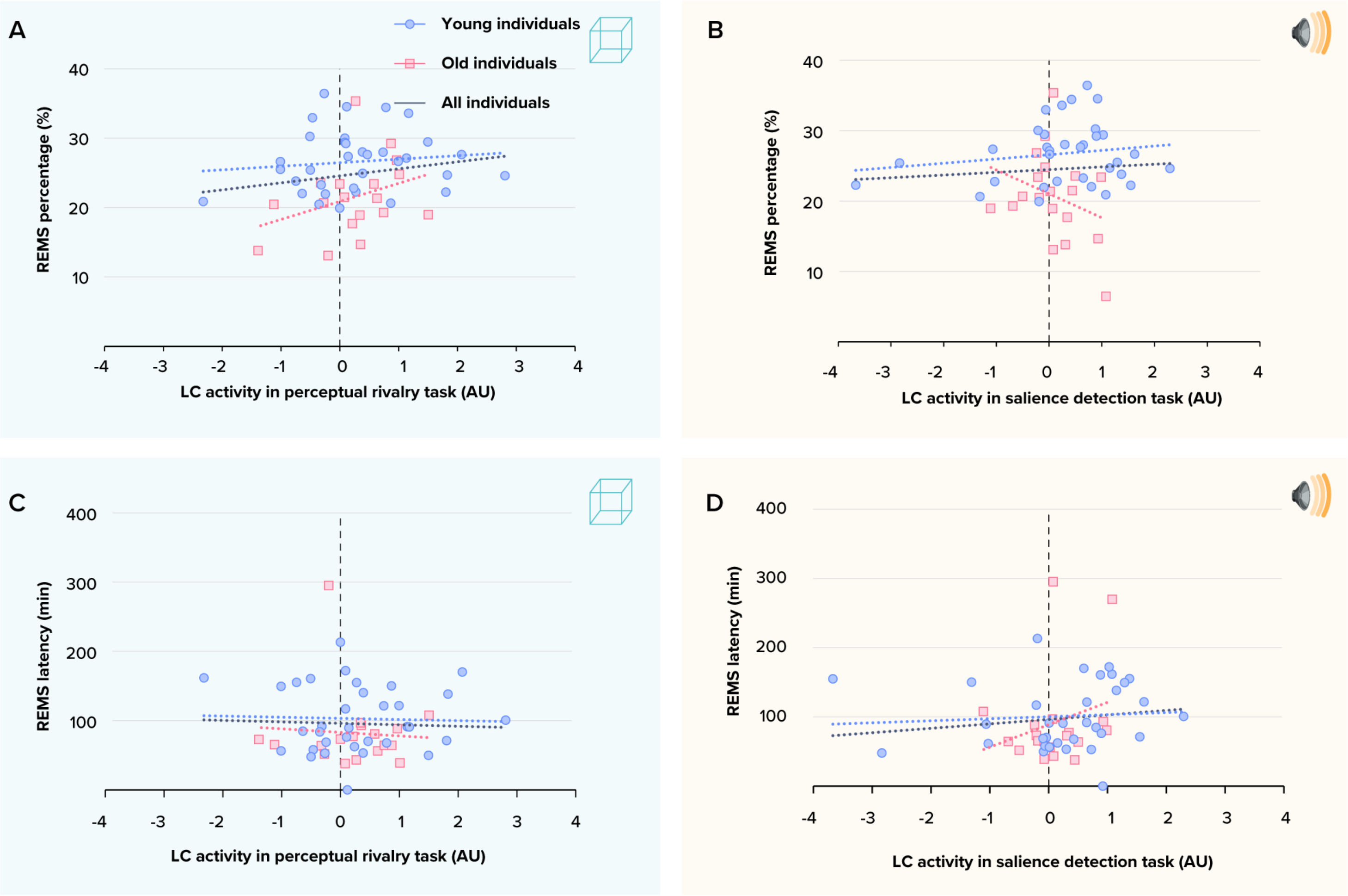


**Supplementary Figure S1. Non-significant associations between the LC activity estimates and main sleep metrics of interest. (A)** Association between REM sleep percentage and the LC activity estimates during the perceptual rivalry task. The GLMM showed neither a significant main effect of LC activity (p=0.159) nor a significant age group by LC activity interaction (p=0.164). **(B)** Association between REM sleep percentage and the LC activity estimates during the salience detection task. The GLMM showed neither a significant main effect of LC activity (p=0.245) nor a significant age group by LC activity interaction (p=0.157). **(C)** Association between REM sleep latency and the LC activity estimates during the perceptual rivalry task. The GLMM showed neither a significant main effect of LC activity (p=0.750) nor a significant age group by LC activity interaction (p=0.816). **(D)** Association between REM sleep latency and the LC activity estimates during the salience detection task. The GLMM showed neither a significant main effect of LC activity (p=0.218) nor a significant age group by LC activity interaction (p=0.242).

Simple regression lines are used for a visual display and do not substitute the GLMM outputs. The black line represents the regression irrespective of age groups (young + old, n = 52). Solid and dashed regression lines represent significant and non-significant outputs of the GLMM, respectively.

155

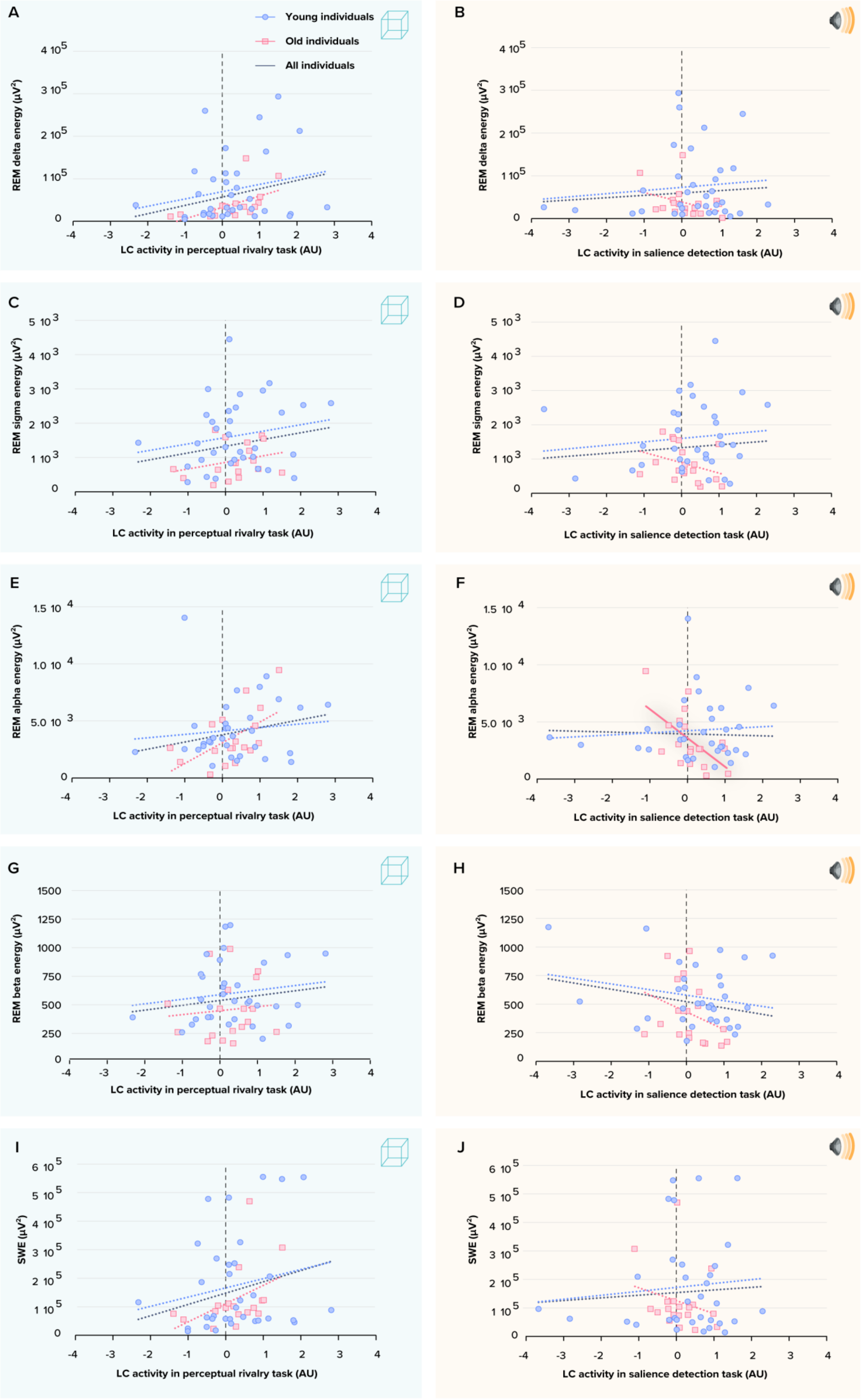
A
Young individuals
Old individuals
All individuals
REM delta energy (µV²)
LC activity in perceptual rivalry task (AU)
B
REM delta energy (µV²)
LC activity in salience detection task (AU)
C
REM sigma energy (µV²)
LC activity in perceptual rivalry task (AU)
D
REM sigma energy (µV²)
LC activity in salience detection task (AU)
E
REM alpha energy (µV²)
LC activity in perceptual rivalry task (AU)
F
REM alpha energy (µV²)
LC activity in salience detection task (AU)
G
REM beta energy (µV²)
LC activity in perceptual rivalry task (AU)
H
REM beta energy (µV²)
LC activity in salience detection task (AU)
I
SWE (µV²)
LC activity in perceptual rivalry task (AU)
J
SWE (µV²)
LC activity in salience detection task (AU)

**Supplementary Figure S2. Associations between the LC activity estimates and sleep metrics of interest to test the specificity.** **(A)** Association between REM delta energy and the LC activity estimates during the perceptual rivalry task. **(B)** Association between REM delta energy and the LC activity estimates during the salience detection task. **(C)** Association between REM sigma energy and the LC activity estimates during the perceptual rivalry task. **(D)** Association between REM sigma energy and the LC activity estimates during the salience detection task. **(E)** Association between REM alpha energy and the LC activity estimates during the perceptual rivalry task. **(F)** Association between REM alpha energy and the LC activity estimates during the salience detection task. **(G)** Association between REM beta energy and the LC activity estimates during the perceptual rivalry task. **(H)** Association between REM beta energy and the LC activity estimates during the salience detection task. **(I)** Association between NREM slow wave energy and the LC activity estimates during the perceptual rivalry task. **(J)** Association between NREM slow wave energy and the LC activity estimates during the salience detection task.

None of the associations were significant ($p>0.057$) except for a significant main effect of LC activity during the salience detection task ($p=.025$) and age-group by LC activity interaction ($p=.037$) when using REM alpha energy as the dependent variable.

Simple regression lines are used for a visual display and do not substitute the GLMM outputs. The black line represents the regression irrespective of age groups (young + old, n = 52). Solid and dashed regression lines represent significant and non-significant outputs of the GLMM, respectively.

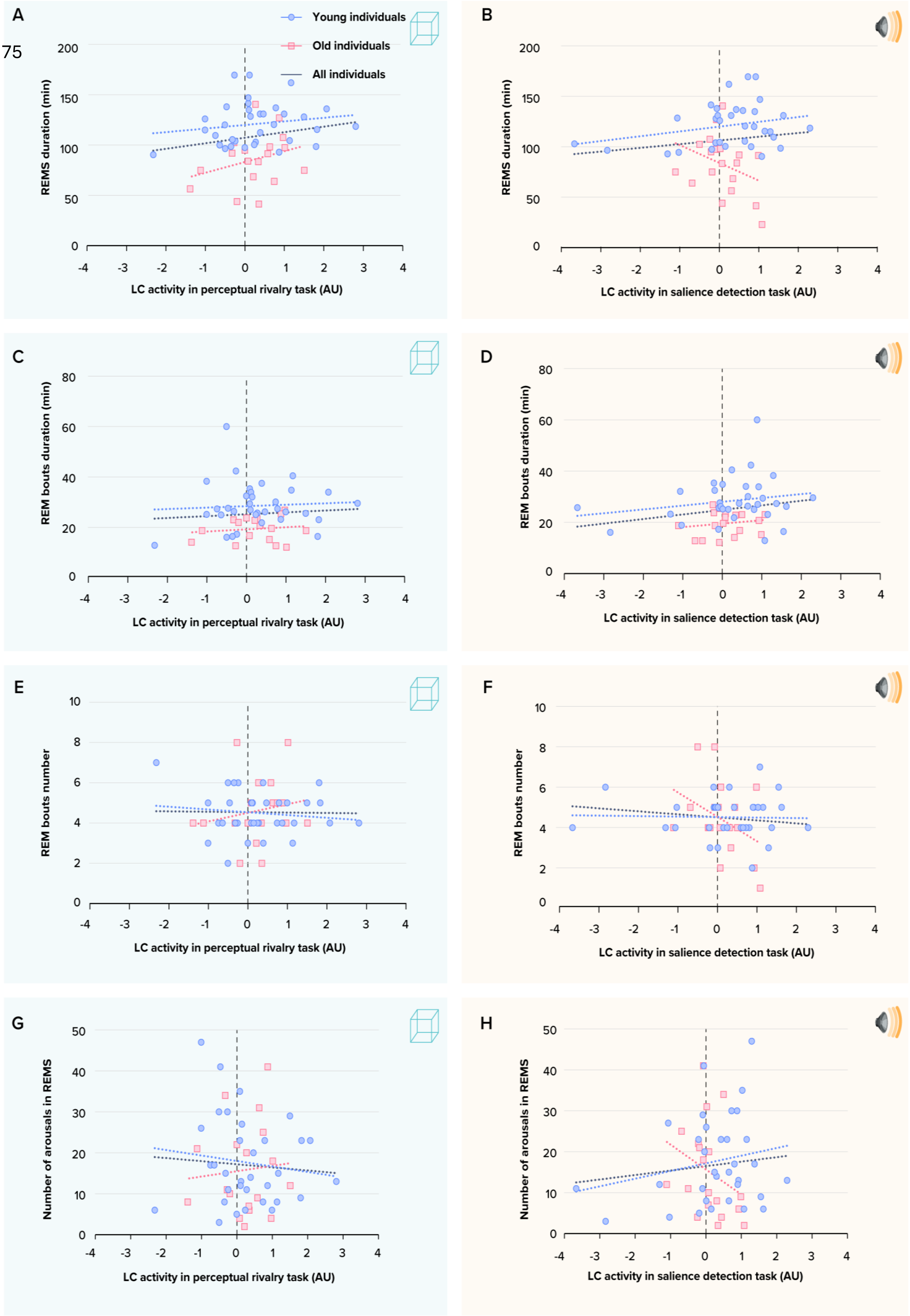
A
Young individuals
Old individuals
All individuals
REMS duration (min)
LC activity in perceptual rivalry task (AU)
B
REMS duration (min)
LC activity in salience detection task (AU)
C
REM bouts duration (min)
LC activity in perceptual rivalry task (AU)
D
REM bouts duration (min)
LC activity in salience detection task (AU)
E
REM bouts number
LC activity in perceptual rivalry task (AU)
F
REM bouts number
LC activity in salience detection task (AU)
G
Number of arousals in REMS
LC activity in perceptual rivalry task (AU)
H
Number of arousals in REMS
LC activity in salience detection task (AU)

**Supplementary Figure S3. Associations between the LC activity estimates and sleep metrics of interest for exploratory analysis. (A)** Association between REM sleep duration and the LC activity estimates during the perceptual rivalry task. **(B)** Association between REM sleep duration and the LC activity estimates during the salience detection task. **(C)** Association between REM bouts duration and the LC activity estimates during the perceptual rivalry task. **(D)** Association between REM bouts duration and the LC activity estimates during the salience detection task. **(E)** Association between REM bouts number and the LC activity estimates during the perceptual rivalry task. **(F)** Association between REM bouts number and the LC activity estimates during the salience detection task. **(G)** Association between number of arousals in REM sleep and the LC activity estimates during the perceptual rivalry task. **(H)** Association between number of arousals in REM sleep and the LC activity estimates during the salience detection task.

None of the associations were significant ($P>0.141$).

Simple regression lines are used for a visual display and do not substitute the GLMM outputs. The black line represents the regression irrespective of age groups (young + old, n = 52). Solid and dashed regression lines represent significant and non-significant outputs of the GLMM, respectively.

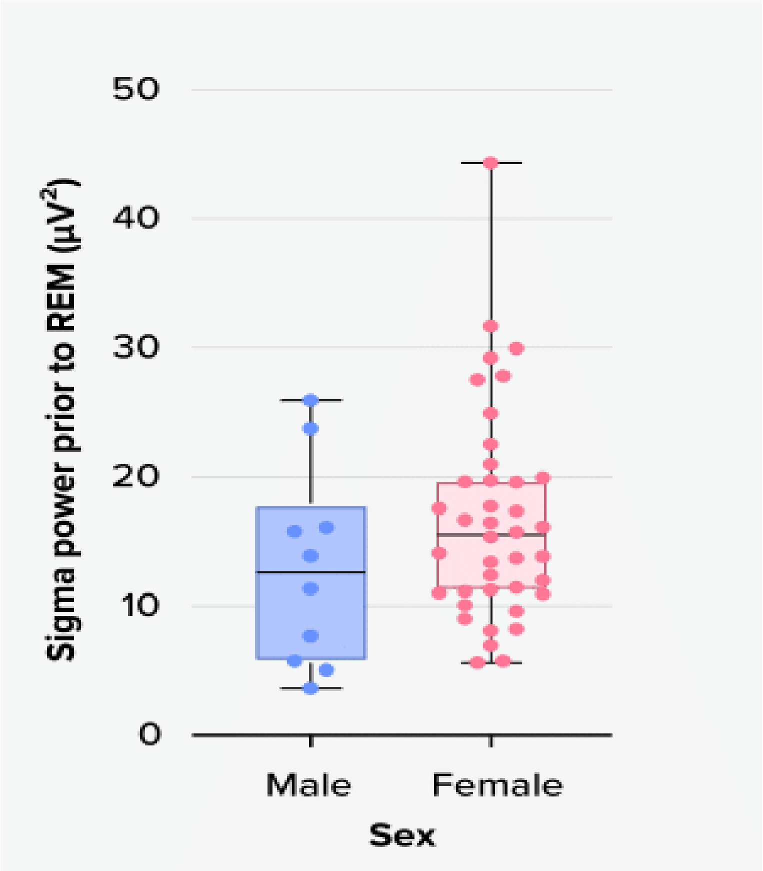


**Supplementary Figure S4. Sigma power prior to REM sleep in males and females.** Men have significantly less sigma power prior to REM sleep compared to women

9. Koshmanova E, Berger A, Beckers E, Campbell I, Mortazavi N, Sharifpour R, et al. Locus coeruleus activity while awake is associated with REM sleep quality in older individuals. JCI

insight. 2023;8.

10. Campbell I, Beckers E, Sharifpour R, Berger A, Paparella I, Aizpurua JFB, et al. Impact of light on task-evoked pupil responses during cognitive tasks. J Sleep Res. 2023;e14101.
